# Microglia are dispensable for experience-dependent refinement of visual circuitry

**DOI:** 10.1101/2023.10.17.562708

**Authors:** Thomas C. Brown, Emily C. Crouse, Cecilia A. Attaway, Dana K. Oakes, Sarah W. Minton, Bart G. Borghuis, Aaron W. McGee

## Abstract

Microglia are proposed to be critical for the refinement of developing neural circuitry. However, evidence identifying specific roles for microglia has been limited and often indirect. Here we examined whether microglia are required for the experience-dependent refinement of visual circuitry and visual function during development. We ablated microglia by administering the colony-stimulating factor 1 receptor (CSF1R) inhibitor PLX5622, and then examined the consequences for retinal function, receptive field tuning of neurons in primary visual cortex (V1), visual acuity, and experience-dependent plasticity in visual circuitry. Eradicating microglia by treating mice with PLX5622 beginning at postnatal day (P) 14 did not alter visual response properties of retinal ganglion cells examined three or more weeks later. Mice treated with PLX5622 from P14 lacked more than 95% of microglia in V1 by P18, prior to the opening of the critical period. Despite the absence of microglia, the receptive field tuning properties of neurons in V1 were normal at P32. Similarly, eradicating microglia did not affect the maturation of visual acuity. Mice treated with PLX5622 displayed typical ocular dominance plasticity in response to brief monocular deprivation. Thus, none of these principal measurements of visual circuit development and function detectibly differed in the absence of microglia. We conclude that microglia are dispensable for experience-dependent refinement of visual circuitry. These findings challenge the proposed critical role of microglia in refining neural circuitry.

## Introduction

Microglia are proposed to be critical mediators of synaptic refinement and neural circuit function^1^. Microglia have been reported to perform essential synaptic pruning in visual circuitry and hippocampus, to mediate motor learning and social interactions, and to contribute to multiple types of circuit refinements^1^. However, data demonstrating that microglia alter neuronal function or corresponding neural circuits relevant to behavior remain limited.

The visual system provides several advantages for quantifying the contribution of microglia to the development, refinement, and plasticity of neural circuits. The function of visual system can be readily measured at the cellular, circuit, and behavior level. The aggregate function of the rod system and cone system can be evaluated with the full field electroretinogram (ERG), and the response properties of retinal ganglion cells can measured with two-photon fluorescence calcium imaging in the *ex vivo* retinal whole-mount retina preparation^2^. Neurons in visual cortex can be characterized for numerous functional properties with calcium imaging, including orientation tuning, spatial frequency (SF) tuning, and binocularity ^3–6^. Each of these tuning properties is generally considered to emerge in visual cortex, although recent studies have detected similar tuning also for some neurons in the visual thalamus, which signals feed-forward visual information to the cortex^7–9^. The visual water task is a validated behavioral assay for quantifying spatial acuity^10^. Acuity measured in this learned behavior assay relies on visual cortex^11^.

The maturation of visual circuitry relies on both activity-dependent refinement and experience-dependent plasticity. In the retina, circuit anatomical structure including lamination of the outer and inner plexiform layers and cell-type specific connectivity is formed before eye opening and prior to visual experience. This circuitry is refined following eye-opening in a stimulus-dependent manner. Refinement not only changes the strength of established synaptic connections, but also eliminates some synaptic connections and adds new ones. Mice lacking the gene for the fractalkine receptor (Cx3cr1), a receptor expressed by microglia in the CNS, are reported to impaired cone-related function^12^.

Retinal ganglion cells project to numerous subcortical nuclei. The major projection for image-forming vision is to the dorsal lateral geniculate nucleus (dLGN) of the thalamus^13^. Retinogeniculate projections are patterned prior to eye opening by spontaneous activity in the form of retinal waves^14,15^. Afferents from the contralateral eye and ipsilateral eye are initially intermingled upon their arrival at the dLGN, but then segregate into separate termination zones with minimal overlap by P14 in the mouse^16,17^. This occurs around the age of natural eye opening. However, mice lacking functional genes for components of the classical complement cascade are reported to display impaired retinogeniculate segregation^18,19^. This defect in activity-dependent refinement is proposed to be a consequence of impaired synapse elimination by microglia ^20^.

The major impact of vision on the development of visual circuitry occurs in primary visual cortex (V1) where the functional properties of neurons can be quantified as specific tuning properties. In mouse, vision drives the refinement of receptive field tuning properties during the first few weeks after eye opening in V1^21–24^. Mean preferred SF and orientation selectivity of excitatory neurons in L2/3 significantly increase between P18 and P36, as does the fraction of binocular neurons^22^. The matching orientation preference for binocular neurons also improves during this period of development^4,6,22^. The refinement of tuning properties accompanies the maturation of visual acuity^25^. This interval during development also overlaps with the ‘critical period’ for the heightened sensitivity of binocularity to abnormal vision. Closing the eye contralateral to the hemisphere of study by lid suture (monocular deprivation, MD) for only 4-5 days between P21 and P32 but not thereafter, shifts eye dominance of neurons in V1 towards the non-deprived (ipsilateral) eye^26^. If microglia perform functions essential for the refinement of synaptic connectivity, then the tuning properties of neurons and their plasticity during the major experience-dependent phase of visual circuit development should be disrupted by their absence.

To eliminate microglia from the developing brain, we fed dams and pups chow containing the small molecule compound PLX5622 (1200mg/kg) that eradicates microglia^27^. Microglia require signaling through the colony-stimulating factor 1 receptor (CSF1R) for survival^28^. PLX5622 is a highly selective, potent, orally bioavailable CSF1R inhibitor that crosses the blood-brain barrier. Treatment can be maintained for durations lasting months^27^.

## Results

First, we performed experiments to test for changes in visual circuitry at the level of the retina following eradication of microglia with the CSF1R inhibitor, PLX5622. The rationale for these experiments was that any change there would necessarily impact all downstream signaling in the visual pathways. To avoid the potential impact of deleting microglia on retinogeniculate segregation, we fed mice chow containing PLX5622 (PLX chow), beginning at P14 concomitant with natural eye opening and after retinogeniculate segregation.

Confocal fluorescence imaging 21 days (d) after feeding mice chow containing PLX5622 (1200mg/kg, PLX chow) revealed a complete absence of microglia at all levels of the retina, consistent with published studies^29^ (Fig. 1a,b). To test for changes at the level of the photoreceptor and ON bipolar cell level, we compared the ERGs of control vs. 35d PLX-treated mice that express enhanced green fluorescent protein (eGFP) from the CX3CR1 gene locus^30^. We did not observe any significant differences in the scotopic a- and b-wave, which represents the rod photoreceptor and rod bipolar cell response (Fig. 1c). We also found no significant difference in the photopic a- and b-wave, which represent the cone photoreceptor and cone ON bipolar cell response, respectively (Fig 1c). Two-photon fluorescence calcium imaging of retinal ganglion cell populations in Thy1-GCaMP6f mice showed no significant differences in response amplitudes (normalized change in fluorescence, delta F/F) ON, OFF, and ON-OFF type responses between the microglia-free and control groups (one-way ANOVA and Holm-Sidak’s multiple comparison test) (Fig. 1d-f)^31^. The frequency distribution of response types was also similar. We conclude that loss of microglia does not cause evident changes in visual signaling in the retina as we find no detectable change in the gross retinal output. This finding is consistent with a previous study that reported no short-term effect of microglial ablation on retinal structure and visual function, but did detect a decrease in the scotopic rod response amplitude^32^.

**Figure 1.**
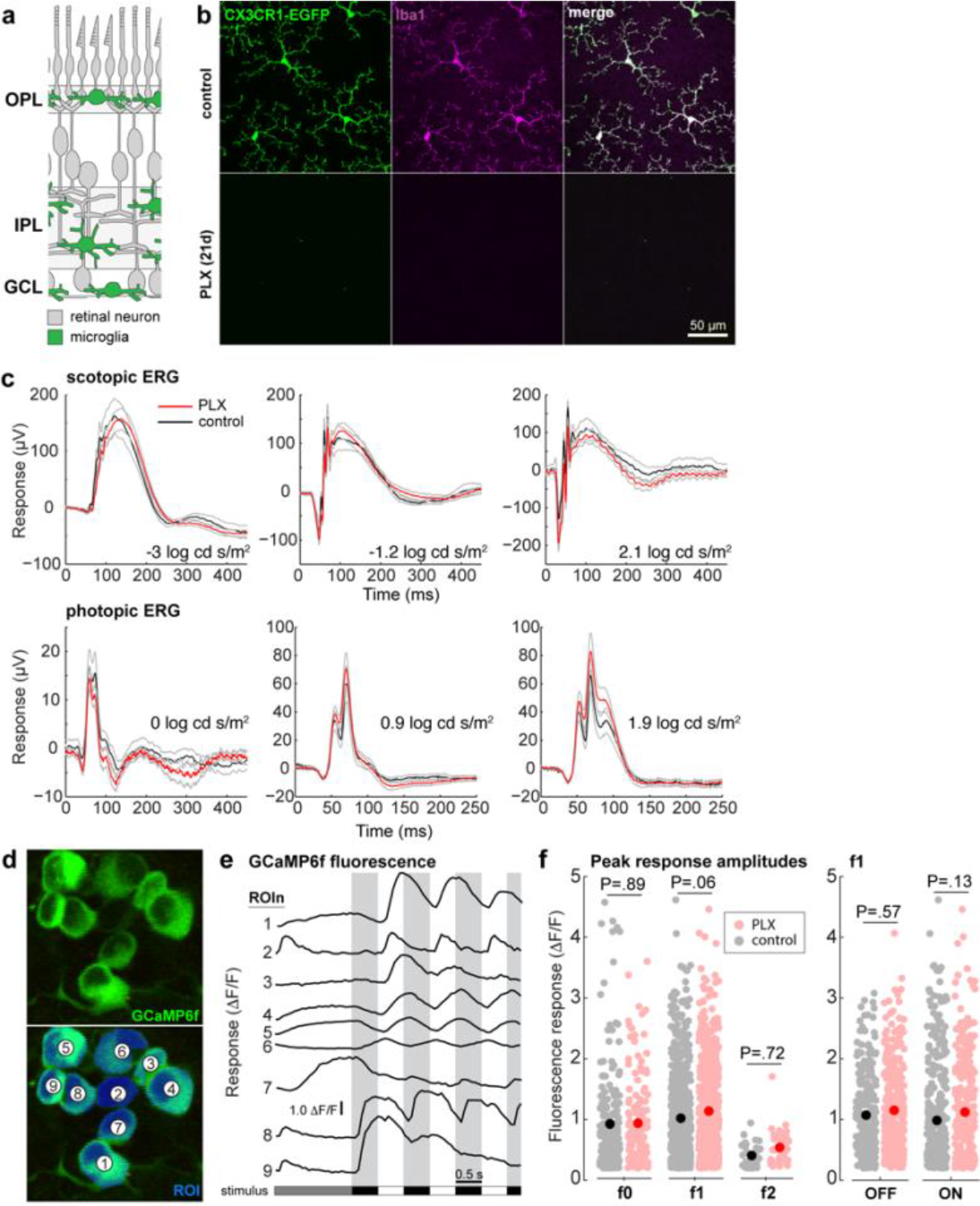
Visual signaling in retina of mice without microglia. **(a)** Schematic distribution of microglia (green) within the retinal anatomy (gray). (**b)** Confocal fluorescence images at the level of the outer plexiform layer (OPL) of control (top) and PLX5622-treated retinas (bottom) of CX3CR1-EGFP mice (green, left) with immuno-histochemical labeling against microglia marker Iba-1 (magenta, center) and the merged image (right). Scale bar = 50 microns. (**c)** Dark adapted (top) and light adapted (bottom) full field electroretinogram with light flashes at three intensities. Black, control; red, PLX-treated; gray lines ± 1.0 S.E.M. (**d)** Two-photon fluorescence image of GCaMP6f-expressing ganglion cells. Bottom image shows regions of interest (ROIs) masks (blue) used to extract the stimulus-evoked calcium response for each cell. **(e)** GCaMP6f fluorescence responses of the cells shown in **d**. Grey bars indicate the dark phase of the 1Hz, contrast-reversing spot stimulus. **(f)**, response amplitudes for all cells recorded in control (gray; mean ± 1.0 SEM, black) and PLX-treated mice (pink; mean ± 1.0 SEM, red). Left panel shows cells grouped by primary response waveform: non-modulating (f0; control n = 189; PLX n = 140), modulating at the stimulus frequency (f1; control n = 565 ; PLX n = 582), and modulating at twice the stimulus frequency (f2; ON-OFF response; control n = 35 ; PLX n = 38). Right panel, response amplitudes of OFF- and ON-type ganglion cells with peak response at the stimulus frequency (1Hz; f1 population of left panel; OFF, control n = 234; PLX n = 309; ON, control n = 331; PLX n = 273).

Eradication of microglia was nearly complete after four days of treatment will PLX5622. In primary visual cortex (V1), only a handful of remaining microglia were evident at P18 by staining against Ionized calcium binding adaptor molecule 1 (Iba-1), an established marker for microglia (Fig. 2a). Less than 1% of the number of microglia present in control mice were evident 7 days after onset of treatment (Fig 2b). This depletion of microglia is consistent with that observed in other studies that delivered chow to weaned mice at P18 and older ages^27,33^.

**Figure 2.**
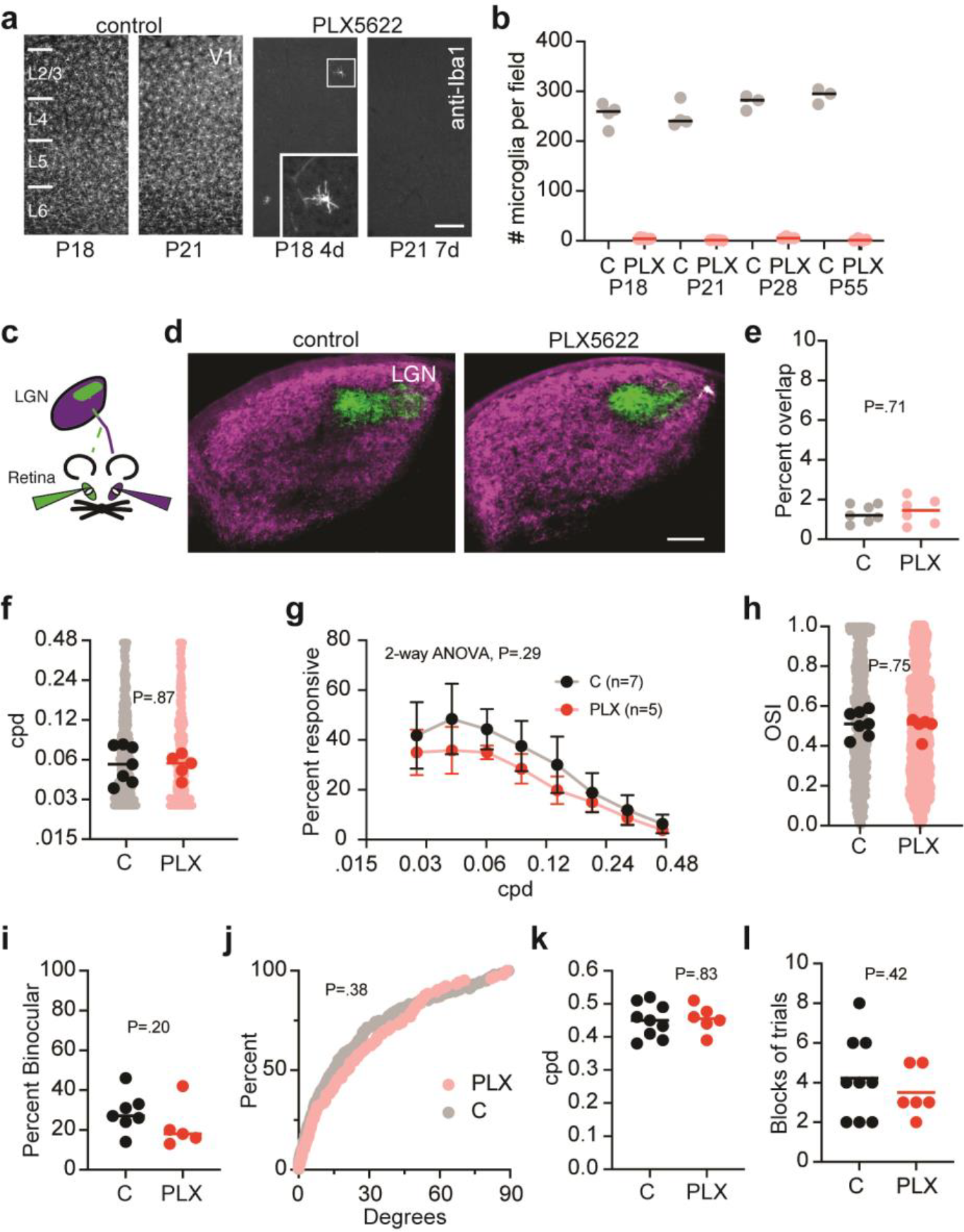
Retinogeniculate segregation in LGN, neuronal tuning properties in V1, and visual acuity of mice without microglia. **(a)** Fluorescence images of primary visual cortex (V1) from control (left) and PLX5622-treated (right) mice at P18 and P21 stained with antibodies directed against Iba-1. Scale bar is 125 microns Inset is 2.5X magnification. **(b)** Number of microglia per imaging shown in panel **a**. Each symbol represents one imaging field. Grey circles are controls (C) and pink circles are PLX-treated (PLX). (P18, C n = 4; PLX n = 8; P21, C n = 4; PLX n = 5; P28, C n = 3; PLX n = 4; P55, C n = 3; PLX n= 4). **(c)** Schematic of the visual pathway from the retinas to LGN and experimental design of the anterograde dual color cholera toxin b. Injections into the eye ipsilateral to the hemisphere of study (green) target a small patch in the LGN surrounded by a larger contralateral (magenta) projection. **(d)** Example images of the overlap of retinogeniculate axons in the LGN as in **c** for control and PLX5622-treated mice. Scale bar = 100 microns. **(e)** Percent overlap of retinogeniculate axons in LGN for control mice (C, grey circles) (n= 7 hemispheres from 5 mice) and PLX-treated mice (PLX, pink circles) (n = 6 hemispheres from 3 mice). **(f)** Preferred spatial frequency (SF) for excitatory L2/3 neurons in V1 from control (C) and PLX-treated mice (C, n = 598; PLX, n = 550) for neurons responsive to the contralateral eye. Black and red circles represent median preferred SF per mouse (C, n = 7; PLX, n = 5). Grey and pink circles represent individual neurons. The lines indicate the median for mice. **(g)** The percent of visually-responsive neurons per mouse that displayed a significant response to each SF at any orientation for control (C) and PLX-treated mice (C, n = 7; PLX, n = 5). **(h)** Orientation selective index scores for the neurons and mice in panel **f. (i)** The percent binocular neurons for mice in panel **f. (j)** The difference in preferred orientation for binocular neurons for mice in panel f (C, n = 203; PLX, n = 104). **(k)** The visual acuity for control (C) and PLX-treated mice measured with the visual water task (C, n = 9; PLX, n = 6). **(l)** The number of blocks of ten trials to reach the testing criteria for each mouse in panel **k**.

We examined whether loss of microglia after eye opening disrupted the established segregation of retinogeniculate inputs to the dLGN. We mapped the distribution of retinal inputs to the dLGN by anterograde tracing with cholera toxin subunit B (CTB). We injected CTB coupled to a different colored fluorophore into each eye on P24 for mice raised from P14 on either control chow or PLX chow (Fig. 2c). Inputs to each hemisphere were dominated by the contralateral eye and surrounded a smaller patch represented by the ipsilateral eye, matching the known normal anatomy (Fig. 2d). Consistent with previous studies, the overlap of territory occupied by inputs from both eyes was small in control mice, just 1-2% (Fig. 2e)^17^. Deleting microglia did not alter this pattern, as the distribution of overlap between to the two eyes was not different from control (C) mice (mean ± SD, C 1.3±.4% vs PLX 1.4±.6%, P = .71, Welch’s t-test). Thus, depletion of microglia after eye opening does not degrade the segregation of these inputs at the first stage of central visual processing.

Next, we measured response tuning properties of visually-driven neurons in primary visual cortex with calcium imaging at cellular resolution in alert mice that expressed the genetically-encoded calcium sensor GCaMP6s^6,34,35^. Mice were maintained on control chow (n=7) or PLX chow (n=5) from P14 until imaging between P28-32. The median preferred SF was nearly identical for the two groups of mice (median C, .058 cycles per degree (cpd) vs. PLX, .059 cpd, P=.87, Mann-Whitney (MW) test), as was the fraction of neurons with significant responses at each SF tested (P=.29, 2-way ANOVA) (Fig. 2f,g)^35^. Orientation tuning was nearly identical for mice raised on control chow or PLX5622 chow (Orientation Selectivity Index, median C .51 vs. PLX .51, P=.75, MW test) (Fig. 2h), and the percentage of binocular neurons was similar between control mice and mice lacking microglia (median C 27% vs PLX 18%, P=.20, MW test) (Fig. 2i). Binocular matching orientation preference was also normal in mice fed PLX chow (median C, 15 degrees vs PLX 21 degrees, P=.38, Kolmogorov-Smirnov (KS) test) (Fig. 2j) Overall, there was no difference between the tuning properties for the two groups at the end of the critical period for experience-dependent refinement.

The maturation of neuronal tuning properties accompanies improved vision. In mice, visual acuity measured with a behavioral assay, the visual water task, nearly doubles in the three weeks after weaning (∼P21) to attain adult levels around P40^10,25^. Visual experience is required for this maturation of acuity^36^. Acuity of mice fed PLX chow (n=6) from P14 to the completion of acuity testing (∼P45) was indistinguishable from mice raised on control chow (n=9) (mean±SD C, .45 cpd vs PLX, .45 cpd; P=.83, Welch’s t test) (Fig. 2k). We conclude that acuity is normal in mice lacking microglia from soon after eye opening through the period of experience-dependent maturation of acuity.

Given that we were unable to detect any effect of eradication of microglia on the experience-dependent refinement of visual circuitry with normal vision during the critical period, we then examined whether microglia contributed to experience-dependent plasticity associated with abnormal vision. Brief monocular deprivation during the critical period shifts ocular dominance towards the non-deprived eye in mammals with binocular vision as measured with multi-unit electrophysiologic recordings across all cortical layers^37,38^. In the mouse, 4 days of monocular deprivation (MD) is sufficient to yield the maximal shift in ocular dominance^26^. Non-deprived control mice exhibit a pronounced bias towards responsiveness to visual stimuli presented to the contralateral eye. This contralateral bias results in higher Contralateral Bias Index (CBI) scores near 0.7 (n=6, mean CBI= .65±.02) (Fig 3a). Four days of MD initiated during the zenith of the critical period (P26-P28) reduces the CBI scores for mice to near 0.5 as ocular dominance shifts significantly away from the contralateral (deprived) eye (n=5, mean CBI=.48±.04, P=.0003, ANOVA) (Fig. 3a). Non-deprived mice maintained on PLX chow (n=5) from P14 to P32 had normal ocular dominance and whereas mice receiving 4-days of MD (n=5) displayed typical shifts in CBI scores (PLX CBI=.66±.02, MD PLX CBI=.45±.03, P=.0001, ANOVA) (Fig. 3a). The CBI scores for control and PLX mice following MD were indistinguishable (P=.30, ANOVA). Thus, ocular dominance (OD) plasticity was normal in mice lacking microglia for the duration of the critical period as measured with electrophysiology.

**Figure 3.**
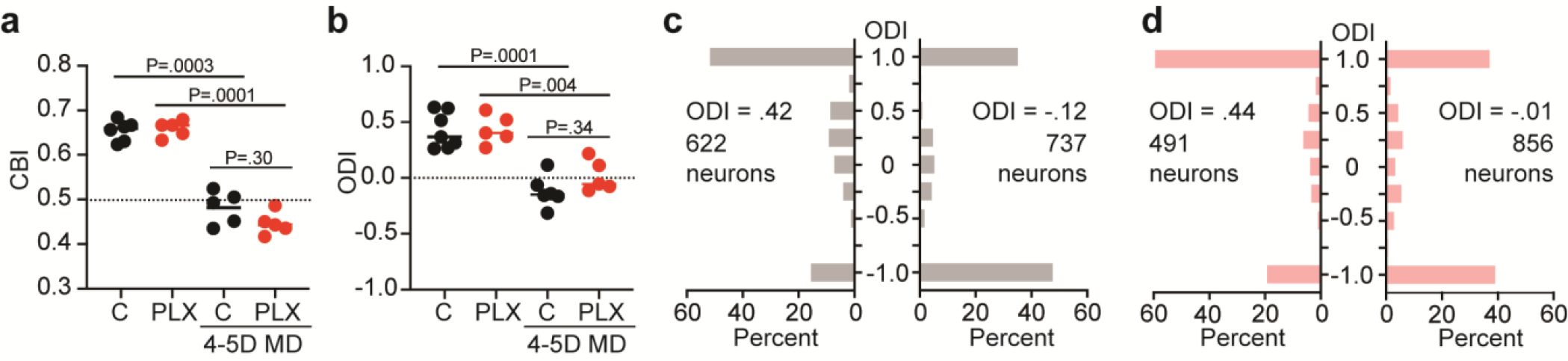
OD plasticity of mice without microglia. **(a)** Contralateral Bias Index (CBI) scores from multi-unit electrophysiologic recordings for non-deprived control (C) mice (n = 6) and non-deprived mice fed PLX chow (PLX) (n = 5), as well as mice receiving 4 days of MD (n = 5 each). **(b)** Ocular Dominance Index scores from calcium imaging for non-deprived control (C) mice (n = 7) and non-deprived mice fed PLX chow (PLX) (n = 5), as well as mice receiving 4 days of MD (n = 6 and 5). **(c)** Histogram of ODI scores for neurons corresponding to the non-deprived and 4d MD control mice in panel **b** (non-deprived, n = 622; MD, n = 737). **(d)** Histogram of ODI scores for neurons corresponding to the non-deprived and 4d MD PLX mice in panel **b** (non-deprived, n = 491; MD, n = 856).

OD plasticity during the critical period involves a complex change in the receptive field tuning properties of excitatory cortical neurons. For example, in layer 2/3, MD reduces the number of excitatory neurons responsive only to the contralateral (deprived) eye and increases the number of neurons responsive only to the ipsilateral (non-deprived) eye^35^. We examined the binocular tuning of neurons in visual cortex of PLX-treated and control mice after 4 days of MD with calcium imaging (Fig. 3b-d). MD yielded a similar OD shift in both groups that was consistent with the OD plasticity detected with multi-unit electrophysiologic recordings (Fig. 3a). Similar to control mice, MD decreased the fraction of neurons that were predominantly monocular and responsive to the contralateral (deprived) eye, and increased the fraction of neurons that were predominantly monocular and responsive to the ipsilateral (non-deprived) eye, with only modest effects on binocular neurons (Fig. 3c,d). Thus, microglia do not appear to contribute to OD plasticity as measured with either multi-unit electrophysiologic recordings or calcium imaging at neuronal resolution.

## Discussion

Microglia are proposed to be critical for the organization and refinement of neural circuits^1,39^. Initial evidence for a role for microglia in the development of neural circuitry was that mice lacking the gene for either *C1q* or *C3* exhibit impaired retinogeniculate segregation. These mutant mice were originally reported to display more than triple the typical overlap of WT mice in projections from the two eyes^40,41^. However, no subsequent studies have identified any deficits in visual processing or vision by *C1q* or *C3* mutant mice. For example, *C1q* mutant mice display normal binocularity and OD plasticity during the critical period^42^. Moreover, recent studies have reported that the deficit in segregation in *C1q* and *C3* mutant mice may be only a small fraction of the original finding at less than a 50% difference^43,44^.

Subsequent studies of hippocampus proposed that microglia are necessary for normal brain development because deletion of *cx3cr1* in mice results in a modest and transient increase in dendritic spine density in by hippocampal CA1 neurons, a partial deficit in long-term potentiation (LTP), and a reduction in the severity of seizures induced by the proconvulsant drug pentylene tetrazole in young (P18) but not adult mice^45^. Microglia have also been proposed play an essential role in motor learning and social interactions^46,47^. However, the circuit modifications accounting for these mostly modest deficits are unclear, although both studies propose defects in synaptic pruning as the underlying mechanism.

Here we tested the model that microglia perform essential functions for the development and refinement of neural circuits by measuring the functional consequences for visual circuitry of eradicating microglia during the period of experience-dependent refinement after eye opening (P14) into early adulthood (∼P60). We did not detect alterations in neuronal tuning properties in the retina, in neuronal tuning properties in visual cortex, in the maturation of visual acuity, or in OD plasticity induced by abnormal visual experience. Thus, if microglia play a role in the refinement and plasticity of visual circuitry, whether by synaptic pruning or another mechanism, it may be restricted to thalamus during early developmental ages and/or without functional significance.

Yet preceding studies have concluded that microglia play a critical role in the development and plasticity of visual circuitry. Microglia have been proposed to be essential for the proper establishment of neural circuits through sensory experience because thalamic neurons cultured *in vitro* from mice lacking a gene upregulated in microglia with visual experience, the TNF family cytokine TWEAK (TNF-associated weak inducer of apoptosis), display a minor increase in the number of ‘bulbous’ dendritic spines^48^. In addition, a study similar in design to the experiments presented here also examined the consequences of treating mice with PLX5622 from P18 to adulthood (P90+)^33^. They reported a decrease in orientation tuning and predicted that mice lacking microglia have impaired visual acuity. In contrast, we did not detect any difference in either orientation tuning of neurons in V1 despite presenting visual stimuli with identical resolution for orientation (30 degree spacing), better resolution for SF (octave vs. half octave), and superior sampling (20 trials vs. 40 trials on average). We also measured visual acuity and it was normal in mice lacking microglia during the period of experience-dependent maturation of acuity.

Microglia have also been proposed to be critical for OD plasticity. Either deletion of the gene for the purinergic receptor P2Y12, a microglial gene, or treatment with clopidogrel, a highly selective antagonist for P2Y12, blocks shifts in eye dominance following MD^49,50^. In these experiments, optical imaging of intrinsic signals was employed to measure OD plasticity. This technique measures changes in reflectance from brain tissue to represent neural activity via the hemodynamic response. It remains unclear how disrupting P2Y12 signaling by microglia would disrupt OD plasticity, but eradicating microglia altogether does not. However, a recent study has reported that disrupting microglial P2Y12 signaling impairs neurovascular coupling, which may have influenced measurements of binocularity inferred from optical imaging of intrinsic signals^51^. A preceding study also examined OD plasticity in mice with PX5622 from P14 to P29. They reported that OD plasticity was absent in mice lacking microglia. Unfortunately, this study was unable to detect the established magnitude of OD plasticity in control mice, and whether these recordings were sufficient for interpretation is in question. In contrast, we measured eye dominance directly with multi-unit electrophysiologic recordings, as well as with calcium imaging at neuronal resolution. Multi-unit recordings are the gold standard for measuring eye dominance and permit comparison across numerous species including cat, primate, and rodent^52–55^ These experiments are identical in design to several of our previous studies^25,56–59^. In our hands, with these two techniques that measure neuronal activity, OD plasticity is normal in mice lacking microglia.

Microglia are reported to selectively remodel developing inhibitory circuits. Treating microglia with PLX5622 by intraperitoneal injection from P1 to P15 has been reported to increase the number of synapses from both parvalbumin-positive and somatostatin positive interneurons onto neurons in L4 of V1 as well as double the frequency of spontaneous inhibitory post-synaptic currents (sIPSCs) by these neurons at P15^39^. Interestingly, this manipulation apparently yields a similar increase in excitatory post-synaptic currents (sEPSCs), such that the balance of excitatory to inhibitory neurotransmission remains unchanged. However, PV interneurons are only evident in V1 beginning at P12 and this population continues to expand until P24^60^. Moreover, the functional maturation of GABAergic inhibition occurs from the time of eye-opening into early adulthood and visually-evoked inhibitory activity is not detected in significant numbers until P19^21,60,61^. We eradicated microglia from P18 onward, during this period of PV neuron maturation, yet the tuning properties of excitatory neurons were unchanged. Likewise, OD plasticity, which requires not only the functional maturation of inhibition, but cortical disinhibition, was normal in the absence of microglia^62,63^. Thus, the magnitude and functional significance of this reported remodeling of inhibitory circuits by microglia remains unclear as it does not appear to affect visual circuitry.

PLX5622 is a specific and potent inhibitor of CSF-1R^27^. The *csf1r* gene is required by osteoclasts, macrophages, and microglia^64^. Constitutive mutant mice for *csf1r* exhibit grossly normal embryonic brain development, but perturbed postnatal brain development, including ventricular enlargement, associated thinning of cerebral cortex, and poor viability^65^. By comparison, mice treated with PLX5622 exclusively during embryogenesis (E3.5 to E15.5) display normal gross brain morphology and only minor sex-specific deficits in hyperactivity and anxiolytic-like behavior as adults^66^. Interestingly, genetic deletion of an enhancer region for the *csf1r* gene abolishes microglia and discrete macrophage populations without perturbing the homeostasis of other macrophages. These mutant mice also exhibit normal brain morphology and do not phenocopy neural defects present in the constitutive null *csf1r* mutant mice^67^. Thus, disrupting CSF-1R signaling in peripheral macrophages at early postnatal ages is the likely cause of the severe brain developmental abnormalities of null *csf1r* mutant mice. We suspect that treatment with PLX5622 age at early postnatal ages (e.g. P1 to P15) may yield similar, but less obvious, disruptions of neural development distinct from any proposed role of microglia in synaptic pruning.

If microglia do indeed prune dendritic spines, the magnitude of this effect is minor and does not yield a detectable change in the tuning properties of neurons in visual cortex, functional vision, or experience-dependent visual plasticity, even when microglia are eradicated for nearly the entire period of experience-dependent refinement from shortly after eye-opening through adulthood. In conclusion, we observe that removing microglia, and thereby the functional roles attributed to microglia including spine pruning, are without evident functional consequences for the experience-dependent phase of visual development. We propose that microglia are dispensable for experience-dependent refinement of visual circuitry.

## Methods

| REAGENT or RESOURCE | SOURCE | IDENTIFIER |
| --- | --- | --- |
| **Deposited data** |  |  |
| Retinal ERG composite data |  | N/A |
| Calculated response amplitudes from RGCs |  | N/A |
| Images of Iba-1 staining of visual cortex |  | N/A |
| Images of retinogeniculate tracing |  | N/A |
| Calculated tuning properties for all neurons from calcium imaging |  | N/A |
| Calculated ocular dominance for all units from multi-unit electrophysiologic recordings |  | N/A |
| **Experimental models: Organisms/strains** |  |  |
| Mouse: B6;DBA-Tg(tedO-GCaMP6s)2Niell/j | The Jackson Laboratory | RRID: ISMR_JAX: 024742 |
| Mouse: B6;Cg-Tg(Camk2a-tTA)1Mmay/DboJ | The Jackson Laboratory | RRID: ISMR_JAX: 007004 |
| Mouse: C57BL/6J-Tg(Thy1-GCaMP6f)GP5.17Dkim/J | The Jackson Laboratory | RRID:IMSR_JAX:025393 |
| B6.129P2(Cg)-Cx3cr1tm1Litt/J | The Jackson Laboratory | RRID:IMSR_JAX:005582 |
| **Software and algorithms** |  |  |
| MATLAB | Mathworks | https://www.mathworks.com/ |
| Processing2 | Processing | https://processing.org/ |
| ImageJ | NIH | RRID:SCR_003070 |
| **Reagents** |  |  |
| Iba-1 antibody Fig.1, Cat# 019-19741 | Wako Chemicals | RRID:AB_839504 |
| Iba-1 antibody Fig. 2, Cat# GTX635363 | Genetex | RRID:AB_2888516 |
| Cholera Toxin Subunit B 488 conjugate, Cat# C22841 | Invitrogen |  |
| Cholera Toxin Subunit B 594 conjugate, Cat# C34777 | Invitrogen |  |

### Lead contact

Further information and requests for resources and reagents should be directed to and will be fulfilled by the Lead Contact, Aaron McGee.

### Data availability

The Deposited data listed in the table above have been deposited in Mendeley data at:

### Experimental model and subjects

All procedures were approved by University of Louisville Institutional Animal Care and Use Committee and were in accord with guidelines set by the US National Institutes of Health. Mice were anesthetized by isoflurane inhalation and euthanized by carbon dioxide asphyxiation or cervical dislocation following deep anesthesia in accordance with approved protocols. Mice were housed in groups of 5 or fewer per cage in a 12/12 light dark cycle. Animals were naive subjects with no prior history of participation in research studies.

### Mice

Retinal experiments were performed on CX3CR1-EGFP (JAX; strain #005582) and Thy1-GCaMP6f mice (JAX; strain #025393). Calcium imaging was performed on mice expressing GCaMP6S in excitatory neurons in forebrain. The *CaMKII-tTA* (stock no. 007004) and *TRE-GCaMP6s* (stock no. 024742) transgenic mouse lines were obtained from Jackson Labs ^34,68^. All other experiments were performed on mice from crosses of *CaMKII-tTA* and *TRE-GCaMP6s* that lacked one of the two transgenes. Mice were genotyped with primer sets suggested by Jackson labs.

### Treatment with PLX5622

PLX5622 was mixed into Openstandard diet with 15kcal% fat (Research Diets) at 1200mg/kg and stored at 4C until use.

### Immunohistology of retina

We tested for the presence of microglia in PLX-treated and control retinas using immunohistochemistry against microglia marker Iba1. Eyes were enucleated and hemisected, and the posterior eye cup including retina placed in 4% PFA (30 mins at 4C) followed by wash to 0.1 M PBS. The fixed retina was removed from the eyecup and incubated with blocking buffer (24 hrs at 4C) followed by incubation with primary antibody against Iba1 (rabbit polyclonal; 1:1000; 3-5 days at 4C) and incubation with fluorescence-tagged secondary antibody (goat-α-rabbit; 1:1000, 24 hrs at 4C). Retinas were radially incised for flat-mounting on a glass microscope slide in Vectashield mounting medium (Vector Laboratories). Images were acquired on an Olympus Fluoview 1200 confocal fluorescence microscope.

### Retinal experiments

Mice receiving PLX chow were examined 35 days after beginning treatment. Gross visual function was assessed via electroretinogram (ERG) as previously described ^69^. Neuronal responses to visual stimulation were assessed using two-photon fluorescence calcium imaging of retinal ganglion cells in Thy-1 GCaMP6f mice as previously described^2^.

### Immunohistology of visual cortex

Sectioning and immunostaining per performed as previously described but with an anti-Iba-1 antibody^25,57^.

### Tracing of retinogeniculate projections and analysis of overlap in dLGN

Mice were perfused transcardially with 4% PFA in PBS and the brains sliced coronally into 70 micron sections with a vibratome. The sections spanning the dLGN were mounted serially. The middle 3 sections of the dLGN for each hemisphere were imaged and analyzed. Epifluorescence microscopy was performed with a BX-51 upright microscope (Olympus) and Retiga EX B 12-bit monochrome camera. Images were through a 10X 0.25NA PLAN objective. Images were down sampled to 8-bit with Photoshop software (Adobe). The brightness of the image was adjusted to fill the full dynamic range. The background was then determined by measuring the mean pixel intensity in a region outside of the dLGN. This value was subtracted for each pixel in the image. A mask was drawn at the circumference of the dLGN. The images were then imported into ImageJ (NIH) and binarized. Each channel was then passed through a threshold set at 0.1 and the images multiplied to yield the pixels overlapping between the two images. The number of overlapping pixels was measured as the numerator for the percentage of overlap. The number of pixels in the unmasked area was measured as the denominator for determining the percentage of overlap.

### Calcium imaging to measure neuronal tuning properties

Imaging and analysis were performed blind to treatment. Methods were identical to our previously published experiments^35^. Wide-field (epifluorescent) and cellular resolution (two-photon laser scanning) imaging experiments were performed though a cranial window. In brief, mice were administered carprofen (5mg/kg) and buprenorphrine (0.1 mg/kg) for analgesia and anesthetized with isoflurane (4% induction, 1-2% maintenance). The hair on the scalp was clipped and mice were mounted on a stereotaxic frame with palate bar and their body temperature maintained at 37° C with a heat pad controlled by feedback from a rectal thermometer (TCAT-2LV, Physitemp). The scalp was resected, the connective tissue removed from the skull, and an aluminum headbar affixed with C&B metabond (Parkell). A circular region of bone 3mm in diameter centered over left visual cortex was removed using a high-speed drill (Foredom). Care was taken to not perturb the dura. A sterile 3mm circular glass coverslip was sealed to the surrounding skull with cyanoacrylate (Pacer technology) and dental acrylic (ortho-jet, Lang Dental). The remaining exposed skull likewise sealed with cyanoacrylate and dental acrylic. Mice recovered on a heating pad. Mice were left to recover for at least 2 days prior to imaging.

After implantation of the cranial window, the binocular zone of visual cortex was identified with wide-field calcium imaging similar to our method for optical imaging of intrinsic signals ^56^. In brief, mice were anesthetized with isoflurane (4% induction), provided a low dose of the sedative chlorprothixene (0.5mg/kg i.p.; C1761, Sigma) and secured by the aluminum headbar. The eyes were lubricated with a thin layer of ophthalmic ointment (Puralube, Dechra Pharmaceuticals). Body temperature was maintained at 37°C with heating pad regulated by a rectal thermometer (TCAT-2LV, Physitemp). Visual stimulus was provided through custom-written software (MATLAB, Mathworks). A monitor was placed 25cm directly in front of the animal and subtended +40 to -40 degrees of visual space in the vertical axis. A horizonal white bar (2 degrees high and 20 degrees wide) centered on the zero-degree azimuth drifted from the top to bottom of the monitor with a period of 8s. The stimulus was repeated 60 times. Cortex was illuminated with blue light (475 ± 30nm) (475/35, Semrock) from a stable light source (intralux dc-1100, Volpi). Fluorescence was captured utilizing a green filter (HQ620/20) attached to a tandem lens (50mm lens, computar) and camera (Manta G-1236B, Allied Vision). The imaging plane was defocused to approximately 200 microns below the pia. Images were captured at 10Hz as images of 1024×1024 pixels and 12-bit depth. Images were binned spatially 4×4 before the magnitude of the response at the stimulus frequency (.125 Hz) was measured by Fourier analysis.

For imaging at cellular resolution, mice were mounted by the headplate atop a spherical treadmill. The monitor was centered on the zero azimuth and elevation 35cm away from the mouse and subtended 45 (vertical) by 80 degrees (horizontal) of visual space. A battery of static sinusoidal gratings were generated in real time with custom software (Processing, MATLAB) as described^6^. Stimulus presentation was synchronized to the imaging data by time stamping the presentation of each visual stimulus to the image acquisition frame number a transistor-transistor logic (TTL) pulse generated with an Arduino at each stimulus transition. Orientation was sampled at equal intervals of 30 degrees from 0 to 150 degrees (6 orientations). Spatial frequency was sampled in 8 steps on a logarithmic scale at half-octaves from 0.028 to 0.48 cycles per degree. An isoluminant grey screen was included (blank) was provided as a 9 ^th^ step in the spatial frequency sampling as a control. Spatial phase was equally sampled at 45-degree intervals from 0 to 315 degrees for each combination of orientation and spatial frequency. Gratings with random combinations of orientation, spatial frequency, and spatial phase were presented at a rate of 4 Hz on a monitor with a refresh rate of 60Hz. Imaging sessions were 10 minutes (2400 presentations in total). Consequently, each combination of orientation and spatial frequency was presented 40 times on average (range 29-56).

Imaging was performed with a resonant scanning two-photon microscope controlled by Scanbox image acquisition and analysis software (Neurolabware). The objective lens was fixed at vertical for all experiments. Fluorescence excitation was provided by a tunable wavelength infrared laser (Ultra II, Coherent) at 920 nm. Images were collected through a 16x water-immersion objected (Nikon, 0.8 NA). Images (512x796 pixels, 520x740 microns) were captured at 15.5 Hz at depths between 150 – 400 microns. Eye movements and changes in pupil size were recorded using a Dalsa Genie M1280 camera (Teledyne Dalsa) fitted with 50 mm 1.8 lens (Computar) and a 800nm long-pass filter (Edmunds Optics). Imaging was performed on alert mice positioned on a spherical treadmill by the aluminum head bar affixed to the skull. The visual stimulus was presented to each eye separately by covering the fellow eye with a small custom occluder.

The imaging series for each eye were motion corrected with the SbxAlign tool. Regions of interest (ROIs) corresponding to excitatory neurons were selected manually with the SbxSegment tool following computation of pixel-wise correlation of fluorescence changes over time from 350 evenly spaced frames (∼4%). ROIs for each experiment were determined by correlated pixels the size similar to that of a neuronal soma. The fluorescence signal for each ROI and the surrounding neuropil were extracted from this segmentation map.

The fluorescence signal for each neuron was extracted by computing the mean of the calcium fluorescence within each ROI and subtracting the median fluorescence from the surrounding perimeter of neuropil ^6,70^. An inferred spike rate (ISR) was estimated from adjusted fluorescence signal with the Vanilla algorithm^71^. A reverse correlation of the ISR to stimulus onset was used to calculate the preferred stimuli ^6,70,72,73^. Neurons that satisfied three criteria were categorized as visually responsive: (1) the ISR was highest with the optimal delay of 4-9 frames following stimulus onset. This delay was determined empirically for this transgenic GCaMP6s mouse ^6^. (2) the SNR was at least one standard deviation greater than spontaneously active neurons. The signal is the mean of the spiking standard deviation at the optical delay between 4-9 frames after stimulus onset and the noise this value at frames -2 to 0 before the stimulus onset or 15-18 after it ^6,72^ (3) and the percent of responses to the preferred stimulus was at least one standard deviation greater than spontaneously active neurons. Visual responsiveness for every neuron was determined independently for each eye. The visual stimulus capturing the preferred orientation and SF was the determined from the matrix of all orientations and SFs presented as the combination with highest average ISR.

The preferred orientation for each neuron was calculated as:

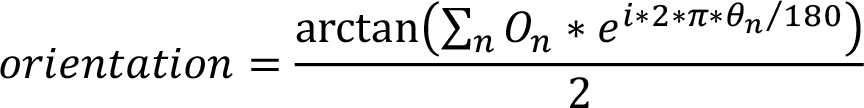

The orientation selectivity index (OSI) was calculated as:

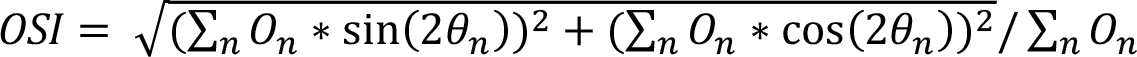

O_n_ is a 1x6 array of the mean z-scores associated with the calculation of the ISR at orientations O_n_ (0 to 150 degrees, spaced every 30 degrees). Orientation calculated with this formula is in radians and was converted to degrees.

The preferred SF for each neuron was calculated as:

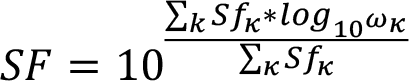

Sf_k_ is a 1x8 array of the mean z-scores at SFs w_k_ (8 equal steps on a logarithmic scale from 0.028 to 0.481 cycles per degree). Tails of the distribution were clipped at 25% of the peak response. The tuning width was the full width at half-maximum of the preferred SF in octaves. The percent visually responsive neurons with significant responses at each SF was determined by comparing the distribution of ISR values at each SF versus the stimulus blank with a KW-test with Dunn’s correction for 8 comparisons. Neurons with P<.01 for a given SF were considered significant responses at that SF.^5^

Binocular matching was measured as the absolute difference in the preferred orientation calculated for visual stimuli presented to the contralateral eye and ipsilateral eye along the 180 degree cycle ^4,23,74^.

Neuronal ODI was calculated as (C-I)/(C+I), where C and I are the mean normalized change in fluorescence (dF/F) for the preferred visual stimulus for the contralateral eye and ipsilateral eye, respectively. In cases where neurons displayed no significant response to visual stimuli provided to one eye, they were considered monocular for the other eye and assigned ODI values of 1 (contralateral) and - 1 (ipsilateral) ^5^. Summed ODI was calculated by summing the dF/F for the preferred visual stimulus for the all neurons visually responsive to the contralateral eye (C) and ipsilateral eye (I) for each mouse, respectively. The summed ODI per mouse was then calculated as (C-I)/(C+I) for each mouse.

### Monocular deprivation

One eye lid was sutured shut on postnatal day 26-28 with 6-0 polypropylene monofilament (Prolene 8709H; Ethicon) under brief isoflurane anesthesia for 4 days. The knot was sealed with cyanoacrylate glue. Upon removing the suture, the eye was examined under a stereomicroscope and mice with scarring of the cornea were eliminated from the study.

### Multi-unit electrophysiologic recordings

Recordings and analysis were performed blind to treatment. Methods were adapted from previously published methods^58,59^. In brief, mice were anesthetized with isoflurane (4% induction, 1-2% maintenance in O_2_ during surgery). The mouse was placed in a stereotaxic frame and body temperature was maintained at 37°C by a homeostatically-regulated heat pad (TCAT-2LV, Physitemp). Dexamethasone (4 mg/kg s.c.; American Reagent) was administered to reduce cerebral edema. The eyes were flushed with saline and the corneas were protected thereafter by covering the eyes throughout the surgical procedure with silicone oil (10838, Millipore Sigma). A craniotomy was made over visual cortex in the left hemisphere and a custom-designed aluminum head bar was attached with Metabond over the right hemisphere to immobilize the animal during recording. Prior to transfer to the recording setup, a dose of chlorprothixene (0.5 mg/kg i.p.; C1761, Sigma) was administered to decrease the level of isoflurane required to maintain anesthesia to 0.8%.

Recordings were made with Epoxylite-coated tungsten microelectrodes with tip resistances of 10 MΩ (FHC). The signal was amplified (model 3600; A-M Systems), low-pass filtered at 3000Hz, high-pass filtered at 300Hz, and digitized (micro1401; Cambridge Electronic Design). Multi-unit activity was recorded from four to six locations separated by >90μm in depth for each electrode penetration. In each mouse, there were four to six penetrations separated by at least 200μm across the binocular region of primary visual cortex, defined by a receptive field azimuth < 25°. Responses were driven by drifting sinusoidal gratings (0.1cpd, 95% contrast), presented in six orientations separated by 30° (custom software, MATLAB). The gratings were presented for 2s of each 4s trial. The grating was presented in each orientation in a pseudorandom order at least four times, interleaved randomly by a blank, which preceded each orientation once. Action potentials (APs) were identified in recorded traces with Spike2 (Cambridge Electronic Design). Only waveforms extending beyond 4 standard deviations above the average noise were included in subsequent analysis. For each unit, the number of APs in response to the grating stimuli was summed and averaged over the number of presentations. If the average number of APs for the grating stimuli was not greater than 50% above the blank, the unit was discarded.

The ocular dominance index (ODI) was calculated as (C-I)/(C+I), where C and I are the average number of APs elicited in a given unit when showing the same visual stimulus to each eye independently. Units were then assigned to one of seven OD categories (1-7) where units assigned to category 1 are largely dominated by input from the contralateral eye, and units assigned to category 7 are largely dominated by input.

## Acknowledgements

This work was supported by a Karl Kirchgessner award (BGB), a grant from the E. Matilda Ziegler Foundation for the Blind (SWM, BGB), a grant from the National Institutes of Health (EY035138 to AWM) and a Jewish Heritage Fund for Excellence Research Enhancement Grant (AWM).

## Author contributions

BGB, AWM, and TCB conceived and designed the study. TCB, ECC, CAA, DKO, SWM, BGB, and AWM performed experiments and analyzed the data. TCB, BGB, and AWM wrote the manuscript.

